# Robust statistical evaluation of tumor incidences in long-term rodent carcinogenicity studies: the reformulated poly-k trend test

**DOI:** 10.1101/2022.02.14.480341

**Authors:** Ludwig A. Hothorn

## Abstract

Mortality-adjusted tumor rates in long-term carcinogenicity rodent bioassays are commonly evaluated by means of the poly-k=3 Armitage trend test. However, this assumes exactly a linear dose-response curve and the Weibull parameter of k=3 for all tumor sites. These unrealistic assumptions can be circumvented by multiple testing across multiple possible dose-response shapes, multiple Weibull parameters, multiple effect sizes, multiple correlated tumors as well as pairwise and trend tests using the multiple marginal models approach. Based on data examples, different multiple tests are demonstrated using the CRAN R packages *multcomp, tukeytrend, coin, MCPAN and multfisher*.

## 1 The problem

In NTP-designed studies, several site-specific tumor incidences in a negative control and a few dose groups are compared for a possible increasing trend, e.g. using the Armitage trend test (CA) [3]. A positive finding is when any tumor type in either males or females *p*^*CA*^ < 0.05- this appears to be a simple and reproducible procedure.

But it is unfortunately not the case: i) tumor development and mortality are interdependent in a complex way, ii) the different tumors found according to an a priori list or spontaneously (and their summaries (e.g., adenomas of the endocrine system) can be modeled as correlated primary endpoints, iii) in some evaluations , pairwise comparisons are performed to control using Fisher tests in addition to the CA-test, iv) the CA test is defined for linear dose-response dependencies (shows highest power there, but not for e.g. plateau-shape profiles), v) either one-or two-tailed trend tests are reported, this has a significant impact on the false negative rate), vi) *p*^*CA*^ < 0.05 vs *p*^*CA*^ < 0.01 relevance criteria for common vs. rare tumors. The specific choice of these six criteria significantly influences the ratio of false positive (f+) to false negative (f−) error rates, the actual and primary issue in these safety studies, where additional effects being discussed. In the following chapter, these criteria are quantified individually from a statistical point of view, with a further underlying issue being various sources of multiplicity. The third underlying problem is the a priori restricted vs. unrestricted formulation of the alternative hypothesis. Based on this balancing of interests, a proposal of an evaluation procedure is made using the multiple marginal tests approach [15] in sub-chapter 2.5.. The idea is not to use only one, a-priori (or per guideline) defined model, but to use several, well-justified models, in order to be able to represent several conceivable data situations. In a maxT test, the best-fitting model then determines the p-value, with the conservativeness price of the multiple models being limited by estimating their correlations. A flexible concept, but with two disadvantages: it is only asymptotically valid, i.e. for infinitely large *n*_*i*_, and so far it does not adequately account for the specific discreteness and sparseness of tumor findings.

## 2 A the reformulated poly-k trend test

### 2.1 Choosing the poly-k parameter k

The relationship between tumor development and mortality is complex: on the one hand, animals that die early cannot develop tumors, on the other hand, tumors can preliminary diagnosed in animals that die or were sacrificed, particularly early. Therefore, the analysis of crude tumor incidences may be biased and should generally be avoided. A simple method of mortality adjustment is the poly-k adjustment [5], where the difficult-to-achieve cause-of-death information in not necessary. Possible individual-specific mortality differences are considered by individual weights *w*_*ij*_ = (*t*_*ij*_/*t*_*max*_)^*k*^ (*t*_*ij*_ … time of death of animal *j* in dose *i*). The weights result in adjusted sample sizes 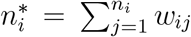 and adjusted proportions 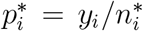 instead of the randomized samples sizes *n*_*i*_ and crude tumor proportions *p*_*i*_ = *y*_*i*_/*n*_*i*_. The shape parameter *k* in a range between *k* = 1, …5 reflects different Weibull hazard functions for cumulative tumor incidences over time. The question arises, which k should be taken specifically (i.e. per tumor site)? The majority of published evaluations take exactly *k* = 3, other studies are performed in parallel for k=3 and k=6 (each to the level *α*). Here we propose a joint *maxT*^*mmm*^ test over all values of *k*, yielding the p-value for the best-fitting k value in each case. Since the six models are highly correlated, the penalty of the multiplicity correction appears to be reasonable.

As an example the incidence of skin fibroma in the bioassay of methyleugenol using rats was used [14], available in the package MCPAN [18]. Table 1 reveals the smallest p-value for poly*k* = 5 (in bold), considerably smaller than the commonly used *k* = 3 model (even if it had been chosen a priori alone). The magnitude of multiplicity adjustment is illustrated by comparing the marginal with the adjusted p-value for k=5 of *p* = 0.0157 and *p* = 0.0205, i.e., if one had known a priori that this was the best assumption. The related R-code is available in the Appendix.

**Table 1:**
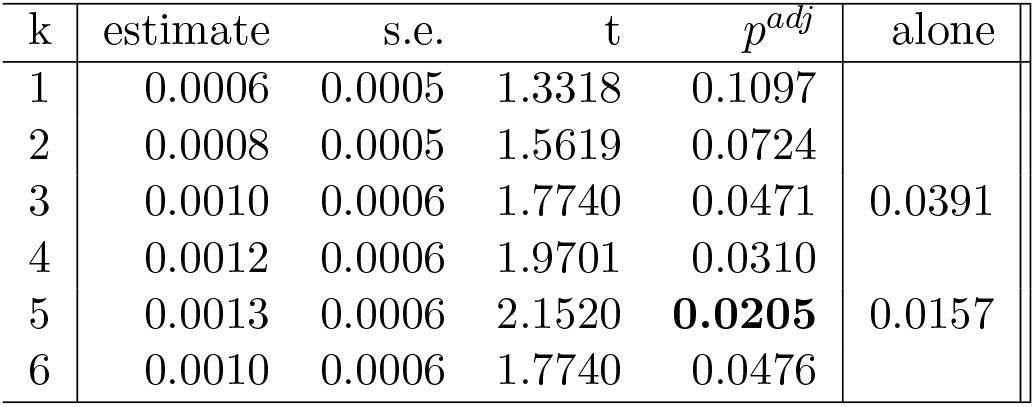
Adjusted p-values for 6 jointly analysed poly-*k* = 1*, …,* 6 tests

This method seems to me recommendable if one imagines that a specific k for the different tumor sites per sex and species is optimal.

### 2.2 Modeling dose as quantitative covariate and qualitative factor, sensitive to more than just linear shapes

The original work of poly-k test [5], is based on the CA trend test [4], formulated precisely for a linear regression model in the generalized linear model (GLM). Given the numerous tumor sites alone, the shape of the dose-response function is not an assumption but a study result. Thus, the use of a trend test is indicated, which guarantees an acceptable power for several shapes, precisely for profiles with plateau shape up to downturns at high dose(s). Already Tukey proposed a maxT test for arithmetic, ordinal and logarithmic (because of D0 precisely arithmetic-logarithmic) dose scores [20], which is now available extended for the glm based on the mmm approach [19]. This approach should be extended by the Williams contrast test [21] (and thus the dose is considered qualitatively and quantitatively, jointly), since this highly-multiple test then detects linear shapes as well as shapes with plateau and even downturns with some power [10]. Although these 4 models are highly correlated, one must accept a multiplicity price. This will be demonstrated by means of the methyleugenol bioassay example (and k=5):

**Table 2:** Tukey-Williams trend test for poly-k=5

| Model | estimate | se | t | $p^{adj}$ |
| --- | --- | --- | --- | --- |
| Regression: arithmetic scores | 0.0013 | 0.0006 | 2.1520 | 0.0357 |
| Regression: ordinal scores | 0.0772 | 0.0272 | 2.8361 | 0.0058 |
| Regression: ari-logarithmic scores | 0.1103 | 0.0392 | 2.8122 | 0.0062 |
| Williams contrast: $p_3 - p_0$ | 0.1640 | 0.0808 | 2.0286 | 0.0472 |
| Williams contrast: $(p_3 + p_2)/2 - p_0$ | 0.1837 | 0.0585 | 3.1377 | 0.0022 |
| Williams contrast: $(p_3 + p_2 + p_1)/3 - p_0$ | 0.1953 | 0.0497 | 3.9303 | 0.0001 |

If one knew a-priori the pooled Williams contrast would be the best-fitting model, then its marginal p-value of only 0.000039 (compared to 0.000109 illustrates the extent of multiplicity- but remember the p-value for an a priori chosen linear CA trend test (k=5) of 0.0157. I.e. the improvement can be substantial, depending on data.

### 2.3 Considering multiple tumor sites simultaneously

One can consider selected or, in extreme cases, all tumor sites as multiple correlated endpoints and thus test them jointly with a maxT test. In clear contrast to issues i and ii), this method is inherently conservative, i.e., a power drop compared to the marginal independent analysis of the individual tumors (each at level *α*) is the consequence. In addition to adherence to the familywise error rate (FWER) (a concept that may not be immediately obvious to toxicologists), the advantage is the joint interpretation of different tumor sites. Once again, the mmm approach is suitable for formulating a double max(max)T test: across multiple tumors and across multiple contrasts or multiple regression models, jointly. Three approaches can be used: i) conditional inference using the package coin [12] (see an example in [7]), ii) multiple contrasts using the package multcomp [11] a further example in [7] and, iii) multiple regression models in the package tukeytrend [19] (see an example in [10]). An example using Williams Trend tests for five selected crude liver tumors (hepatoblastoma, hepatocellular adenoma, hepatocellular carcinoma, hepatocholangiocarcinoma (t24,t27,t29,t30)) in female mice from dataset NTP-TR491 [2] is demonstrated below (see the R-code in the Appendix):

**Table 3:** Multiple tumors Williams trend test

| Tumor site | Contrast | estimate | se | t | p |
| --- | --- | --- | --- | --- | --- |
| Hepatoblastoma | $p_3 - p_0$ | 2.7279 | 1.0620 | 2.5685 | 0.0093 |
| | $(p_3 + p_2)/2 - p_0$ | 2.7279 | 1.0620 | 2.5685 | 0.0093 |
| | $(p_3 + p_2 + p_1)/3 - p_0$ | 2.5089 | 1.0708 | 2.3430 | 0.0168 |
| Hepatocellular adenoma | $p_3 - p_0$ | 2.1160 | 0.4662 | 4.5393 | 0.000004 |
| | $(p_3 + p_2)/2 - p_0$ | 2.3893 | 0.4744 | 5.0364 | 0.0000002 |
| | $(p_3 + p_2 + p_1)/3 - p_0$ | 2.5155 | 0.4788 | 5.2542 | 0.00000008 |
| Hepatocellular carcinoma | $p_3 - p_0$ | 5.0304 | 1.0593 | 4.7487 | 0.000001 |
| | $p_3 - p_0$ | 4.6362 | 1.0537 | 4.4000 | 0.000008 |
| | $(p_3 + p_2 + p_1)/3 - p_0$ | 4.3758 | 1.0515 | 4.1613 | 0.00005 |
| Hepatocholangiocarcinoma | $p_3 - p_0$ | 1.1386 | 1.1719 | 0.9716 | 0.2330 |
| | $p_3 - p_0$ | 0.5693 | 1.2020 | 0.4737 | 0.4119 |
| | $(p_3 + p_2 + p_1)/3 - p_0$ | 0.3795 | 1.1654 | 0.3257 | 0.4713 |

In the joint observation, 4 of the 5 liver tumors show a clear increase in incidences, with the hepatocellular adenoma showing the clearest increase in the shape of a plateau (R-code see in the Appendix).

### 2.4 Choosing effect size: RD, RR, OR

The usual approach is to define the effect size a priori, depending on the design, the test, the interpretation, and so on. When using the p-value, the effect size is often hidden. Mostly the different effect sizes lead to a very different magnitude of the alternative. One can choose some meaningful effect sizes and formulate a maxT- test using the mmm approach [10], e.g. for proportions the risk difference (RD), risk ratio (RR) and odds ratio (OR). When *p*_0_ = 0 is possible or small sample sizes occur, the add-1 adjustment can be recommended [17, 1]. As an example, the poly-k=3 Tukey-type trend tests for hepatoblastoma in female mice [2] is demonstrated below (see the R-code in the Appendix):

**Table 4:** Multiple effect sizes for poly-k=5 Tukey trend tests

| Effect size | Dose score | estimate | se | t | p |
| --- | --- | --- | --- | --- | --- |
| Odds ratio | arithmetic | 0.8458 | 0.2249 | 3.7607 | 0.0003 |
|  | ordinal | 0.8458 | 0.2249 | 3.7607 | 0.0002 |
|  | ari.-logarithmic | 1.5428 | 0.4112 | 3.7516 | 0.0003 |
| Risk difference | arithmetic | 0.1321 | 0.0209 | 6.3313 | < 0.00001 |
|  | ordinal | 0.1321 | 0.0209 | 6.3313 | < 0.00001 |
|  | ari.-logarithmic | 0.2064 | 0.0327 | 6.3140 | < 0.00001 |
| Risk ratio | arithmetic | 0.6215 | 0.1715 | 3.6232 | 0.0004 |
|  | ordinal | 0.6215 | 0.1715 | 3.6232 | 0.0004 |
|  | ari.-logarithmic | 1.1766 | 0.3260 | 3.6089 | 0.0003 |

In this specific example, risk difference as an effect measure shows the lowest p-value.

### 2.5 Pairwise tests vs. trend test

Sometimes trend tests and pairwise tests (control vs. doses) are performed, presumably to be robust against downturns at high doses. If these elementary tests are then performed each at the *α* level, the false positive rate increases significantly. Both principles can also be tested simultaneously to keep the FWER [7]. The first approach based on the conditional inference using the package coin [12] . For the hepatoblastoma in female mice example, the contrasts for Williams and Dunnett test [13] are written explicitly (see the R-code in the Appendix) whereas the first Williams and first Dunnett contrast, namely *p*_3_ − *p*_0_ are double and therefore complete correlated (denoted as double):

**Table 5:**
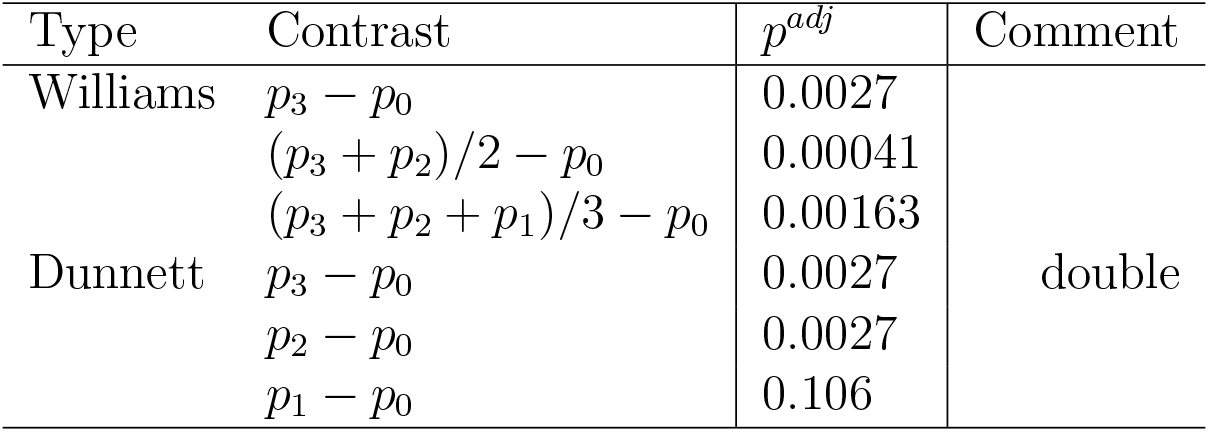
Permutative Dunnett-Williams test for poly-k=5

Not surprisingly in this data example, the smallest p-value for the partial plateau profile of (*p*_3_ + *p*_2_)/2 − *p*_0_. Alternatively, one can take the asymptotic Dunnett-Tukey approach, demonstrated for the skin fibroma example using k=5 (see chapter 2.1):

The magnitude of the increase in the false positive rate can be estimated from the 3 p-values for the Dunnett contrast *p*_1_ − *p*_0_: i) adjusted for both Tukey and Dunnett test *p*^*adj*^ = 0.0065, the Dunnett test alone *p*^*Dunnett*^ = 0.0057 and the unadjusted pairwise contrast *p*^*unadj*^=0.0019 (notice, for asymptotic tests).

**Table 6:**
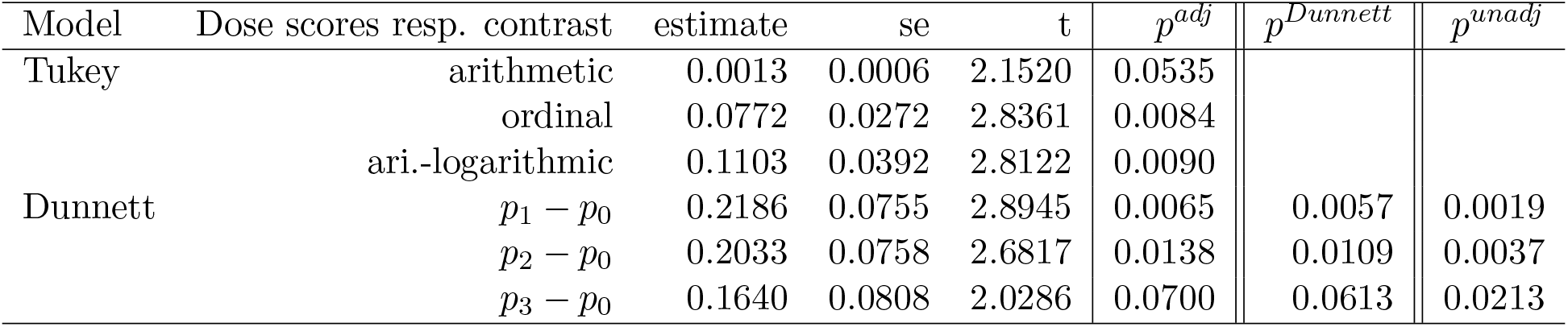
Tukey-Dunnett trend test for poly-k=5

| Model | Dose scores resp. contrast | estimate | se | t | $p^{adj}$ | $p^{Dunnett}$ | $p^{unadj}$ |
| --- | --- | --- | --- | --- | --- | --- | --- |
| Tukey | arithmetic | 0.0013 | 0.0006 | 2.1520 | 0.0535 |  |  |
|  | ordinal | 0.0772 | 0.0272 | 2.8361 | 0.0084 |  |  |
|  | ari.-logarithmic | 0.1103 | 0.0392 | 2.8122 | 0.0090 |  |  |
| Dunnett | $p_1 - p_0$ | 0.2186 | 0.0755 | 2.8945 | 0.0065 | 0.0057 | 0.0019 |
| | $p_2 - p_0$ | 0.2033 | 0.0758 | 2.6817 | 0.0138 | 0.0109 | 0.0037 |
| | $p_3 - p_0$ | 0.1640 | 0.0808 | 2.0286 | 0.0700 | 0.0613 | 0.0213 |

### 2.6 Consideration of covariates

If there could be a relationship between tumor progression and body or organ weights, a covariance analysis in the GLM would be useful. Here, too, several models can be tested simultaneously, e.g. without covariate, with exactly one or all covariates, see e.g. the relative organ weight issue [7].

### 2.7 The impact of two vs. one-tailed tests on f+ rate

Two-sided testing is not problem adequate , as only increasing tumor incidences are of interest and unnecessarily conservative. This is demonstrated using the Tukey-Dunnett test from chapter 2.5):

**Table 7:**
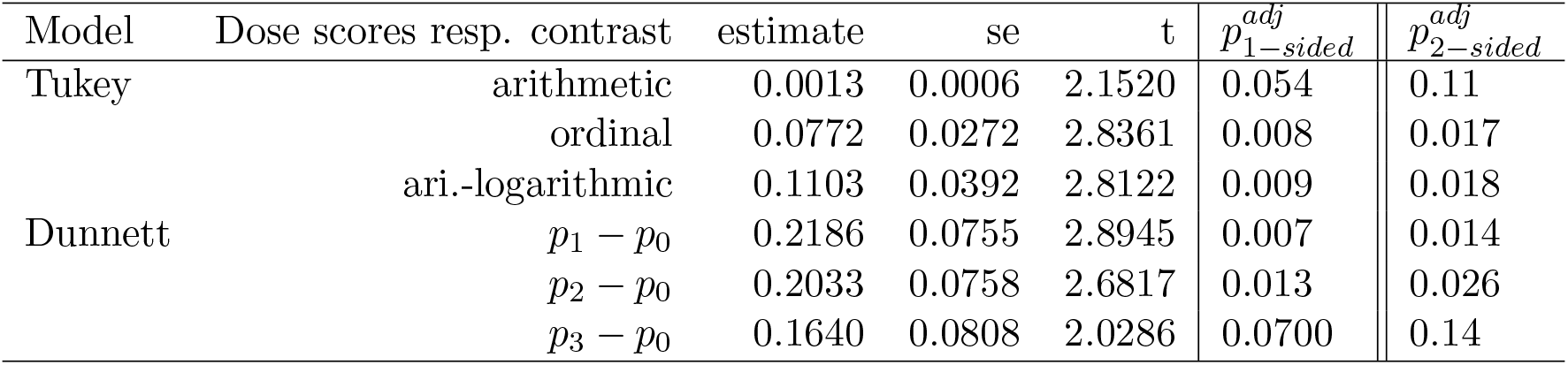
Tukey-Dunnett trend test for poly-k=5: 1-sided vs. 2 sided hypotheses

### 2.8 The availability of simultaneous confidence interval

We have been in a p-value centered interpretation. This has advantages and disadvantages. Therefore, the availability of simultaneous confidence intervals, as compatible as possible with the adjusted p-values, is an advantage - and this is exactly what the mmm-approach offers. As an example the two-sided confidence limits for the six poly-k parameters (*k* = 1, 2, 3, 4, 5, 6) and the three tumor sites (hepatoblastoma, hepatocellular adenoma, hepatocellular carcinoma) can be estimated with the code in the Appendix.

### 2.9 An approach which includes many models simultaneously

In the following, we demonstrate that even multiple models can be considered simultaneously in one approach: i) multiple k, ii) multiple regression models, and iii) multiple tumor sites. Not that I would recommend such a complex approach for routine re-evaluation. The point here is to illustrate the trade-off between the broadest possible alternatives and still permissible conservatism by means of an example using three tumor sites (hepatoblastoma, hepatocellular adenoma, hepatocellular carcinoma; t24,t27,t29), 2 effect sizes (odds ratio, risk difference; OR, RD), 6 poly-k parameter (*k* = 1, 2, 3, 4, 5, 6) and 3 regression models (arithmetic, ordinal, ari.logarithmic), which results in the crazy number of 108 models to be tested simultaneously:

**Table 8:**
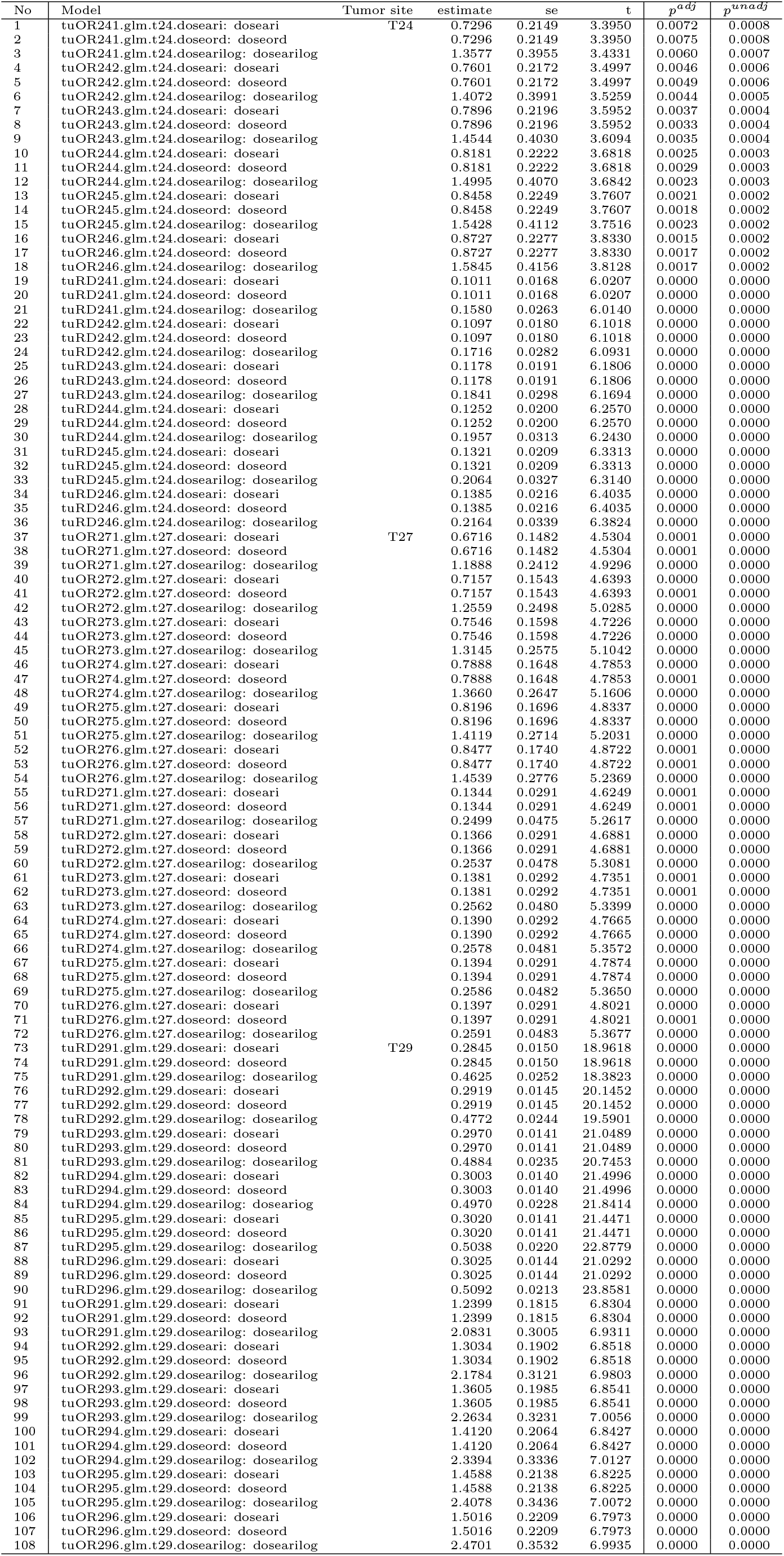
Considering multiple tumors, poly-k parameters, effect sizes and regression models jointly

However, the multiplicity penalty is not too big. E.g., for the hepatocellular adenoma (T27), the standard procedure (poly-k=3 CA test) yields an unadjusted p-value of 0.000054. In comparison, the adjusted p-value of 0.000068 is only slightly higher, compared to the Bonferroni adjusted p-value of 0.0059.

### 2.10 Further modeling aspects

There are several conditions that can be selected, which influence the ratio f+/f−. According to the guidelines, not exactly 1 study per se is evaluated, but several: females and males, rats and mice. The standard is currently to analyze them separately, each at the alpha level. From a statistical point of view, this is not mandatory. For example, one can analyze tumors of the liver for a factorial plant with primary factor dose and secondary factor sex. One could also analyze this for selected tumors across rats and mice in this way. For some important substances, there are even several studies from several laboratories with widely varying dosages. These can be evaluated e.g. in a mixed model [9]. Furthermore, the spontaneous rate of the current control *p*_0_ as well as the historical control has a significant influence - poly-k approaches should be made available for this.

An interesting aspect is the influence of the of unbalanced design on f+ rate. First, from the perspective of duplicate controls, and second, lower *n*_*i*_ due to group-specific mortality patterns [8].

## 3 Taking discreteness into account

In multiple testing of proportions, when using asymptotic tests, the level cannot be exhausted, which occurs due to the discreteness of the data (especially when *p*_0_ = 0). One can use approximations, like add1 [17], or conditional or unconditional tests which use the FWER optimally. Only a few algorithms are currently available for this, and mostly for the two-sample case. With the help of the closed testing procedure, multiple tests can be formulated. In the following, two exact methods are empirically compared with the asymptotic approach on the basis of the mice dataset for the three crude proportions for hepatoblastoma, hepatocellular adenoma, hepatocellular carcinoma.

First, the condition coin-based Dunnett-type test, i.e. a conditional exact max(max)T-test [12], second the exact multivariate Fisher test [16] Bonferroni-adjusted for the 3 two-sample designs *Cvs.D*_*j*_ and the asymptotic add-1 approach [17].

**Table 9:** Crude proportions: exact tests vs. asymptotic test

| Tumor site | Contrast | $p_{coin-\max(\max)}^{adj}$ | $p_{multiFisher}^{adj}$ | $p_{add-1asymptotic}^{adj}$ |
| --- | --- | --- | --- | --- |
| Hepatoblastoma | $p_1 - p_0$ | 0.266 | 0.040 | 0.85 |
| | $p_2 - p_0$ | 0.0067 | 7.51e-10 | 0.48 |
| | $p_3 - p_0$ | 0.0067 | 4.57e-09 | 0.48 |
| Hepatocellular adenoma | $p_1 - p_0$ | 1.39e-09 | 0.00079 | 0.0065 |
| | $p_2 - p_0$ | 5.11e-09 | 2.82e-09 | 0.0088 |
| | $p_3 - p_0$ | 5.72e-06 | 1.60e-10 | 0.046 |
| Hepatocellular carinoma | $p_1 - p_0$ | 6.69e-06 | 0.0024 | 0.022 |
| | $p_2 - p_0$ | 2.41e-08 | 8.46e-09 | 0.0044 |
| | $p_3 - p_0$ | 8.00e-14 | 4.81e-10 | 0.00019 |

On the one hand, the exact estimation of very small p-values is imprecise, on the other hand, from a toxicological point of view it does not matter whether a p-value is 8.00*e* 14 or 4.81*e* 10. But the potential of exact tests in this 2 times k panel data seems to be considerable. However, currently even low-dimensional models are extremely computationally expensive, so much so that routine evaluation is rather difficult.

## 4 Summary

The current standard for evaluating mortality-adjusted tumor rates in long-term carcinogenicity studies is to use the poly-k=3 Armitage trend test, per tumor separately and each at the *α* level. Assuming that a trend holds exactly on a linear alternative and exactly for the Weibull distribution parameter *k* = 3 for all tumor sites seems quite unrealistic. Here, we propose a method of multiple testing based on the multiple marginal models approach with control of FWER that jointly models different dose-response models (where dose is modeled both quantitatively and qualitatively), different Weibull parameters k, different effect sizes, multiple covariates, multiple correlated tumors as well as pairwise and trend tests. Of course, this approach is more conservative than the independent marginal tests, which result in a higher false positive rate. But not extremely conservative (like a Bonferroni test), because the correlations between all marginal tests are taken into account. The basic problem is that a not precisely formulated question (”does a trend exist for this tumor?”) is answered with a multitude of possible alternatives. This is robust, but conservative. This new approach can be evaluated numerically with existing CRAN R packages. The main limitation is that it is defined for asymptotic tests, but the real *n*_*i*_ can be very finite, especially since the table data can be very sparse with the statistically challenging *p*_0_ = 0 case. A way out could be the use of exact tests, which then make exploitation of the level better. A sketch for this has been implemented, but additional work is needed for multiple testing for sparse discrete data, e.g. following [6].

Although the level of complexity outlined above will certainly frighten toxicologists (and is certainly not suitable for routine use to this extent)-the current poly-k=3 Armitage test carries the risk of unacceptably high false negative rates when analyzing one tumor, but also high false positive rates given > 50 tumors considered.

## 5 Appendix: R-code

## 5.1 R-code: poly k=1,…,6 trend tests

~~~
################# all k CA test
library(MCPAN)
data(“methyl”, package=“MCPAN“)
data(methyl)
me <− methyl
me$weightpoly1 <− 1; me$weightpoly2 <− 1; me$weightpoly3 <− 1
me$weightpoly4 <− 1; me$weightpoly5 <− 1; me$weightpoly6 <− 1
# Animals without tumor at time of death get corrected sample size
wt0 <− which(me$tumour == 0)
me$weightpoly1[wt0] <− (me$death[wt0]/max(me$death))^1
me$weightpoly2[wt0] <− (me$death[wt0]/max(me$death))^2
me$weightpoly3[wt0] <− (me$death[wt0]/max(me$death))^3
me$weightpoly4[wt0] <− (me$death[wt0]/max(me$death))^4
me$weightpoly5[wt0] <− (me$death[wt0]/max(me$death))^5
me$weightpoly6[wt0] <− (me$death[wt0]/max(me$death))^6
me$dosegroup <− me$group
levels(me$dosegroup) <− c(“0”, “37”, “75”, “150“)
me$dose <− as.numeric(as.character(me$dosegroup))
# Notice, the modell use an identity link for RD,
fitpoly1 <− glm(tumour ~ dose, data=me, family=binomial(link=“identity“), weight=weightpoly1)
fitpoly2 <− glm(tumour ~ dose, data=me, family=binomial(link=“identity“), weight=weightpoly2)
fitpoly3 <− glm(tumour ~ dose, data=me, family=binomial(link=“identity“), weight=weightpoly3)
fitpoly4 <− glm(tumour ~ dose, data=me, family=binomial(link=“identity“), weight=weightpoly4)
fitpoly5 <− glm(tumour ~ dose, data=me, family=binomial(link=“identity“), weight=weightpoly5)
fitpoly6 <− glm(tumour ~ dose, data=me, family=binomial(link=“identity“), weight=weightpoly6)
library(tukeytrend)
library(multcomp)
ttpoly1 <− tukeytrendfit(fitpoly1, dose=“dose”, scaling=“ari“)
ttpoly2 <− tukeytrendfit(fitpoly2, dose=“dose”, scaling=“ari“)
ttpoly3 <− tukeytrendfit(fitpoly3, dose=“dose”, scaling=“ari“)
ttpoly4 <− tukeytrendfit(fitpoly4, dose=“dose”, scaling=“ari“)
ttpoly5 <− tukeytrendfit(fitpoly5, dose=“dose”, scaling=“ari“)
ttpoly6 <− tukeytrendfit(fitpoly6, dose=“dose”, scaling=“ari“)
compttpoly1 <− glht(model=ttpoly1$mmm, linfct=ttpoly3$mlf, alternative=“greater“)
compttpoly2 <− glht(model=ttpoly2$mmm, linfct=ttpoly3$mlf, alternative=“greater“)
compttpoly3 <− glht(model=ttpoly3$mmm, linfct=ttpoly3$mlf, alternative=“greater“)
compttpoly4 <− glht(model=ttpoly4$mmm, linfct=ttpoly3$mlf, alternative=“greater“)
compttpoly5 <− glht(model=ttpoly5$mmm, linfct=ttpoly3$mlf, alternative=“greater“)
compttpoly6 <− glht(model=ttpoly6$mmm, linfct=ttpoly3$mlf, alternative=“greater“)
tt16 <− combtt(ttpoly1,ttpoly2, ttpoly3,ttpoly4, ttpoly5,ttpoly6)
TT16 <− summary(asglht(tt16, alternative=“greater“))
~~~

## 5.2 R-code: Tukey-Williams poly-k=5 trend tests

~~~
####### k=5 Tukey-Williams test
TW5 <− tukeytrendfit(fitpoly5, dose=“dose”,
scaling=c(“ari”, “ord”, “arilog”, “treat“), ctype=“Williams“)
tw5 <− glht(model=TW5$mmm, linfct=TW5$mlf, alternative=“greater“)
polytW5<−fortify(summary(tw5))$test$pvalue
#### marginal Williams contrast
me$Dose <− as.factor(me$dose)
fitpoly5a <− glm(tumour ~ Dose, data=me, family=binomial(link=“identity“), weight=weightpoly5)
TWi5<−summary(glht(fitpoly5a,linfct= mcp(Dose=“Williams“), alternative=“greater“), adjusted(“none“))
~~~

## 5.3 R-code: Williams trend tests for five correlated liver tumors

~~~
library(multcomp)
glm24 <− glm(cbind(Successes + 1, 52 - (Successes+1)) ~ Group,
subset(miceN, Site==“t24“), family=binomial())
glm27 <− glm(cbind(Successes + 1, 52 - (Successes+1)) ~ Group,
subset(miceN, Site==“t27“), family=binomial())
glm29 <− glm(cbind(Successes + 1, 52 - (Successes+1)) ~ Group,
subset(miceN, Site==“t29“), family=binomial())
glm30 <− glm(cbind(Successes + 1, 52 - (Successes+1)) ~ Group,
subset(miceN, Site==“t30“), family=binomial())
mTW <−summary(glht(mmm(T24=glm24,T27=glm27,T29=glm29,T30=glm30),
mlf(mcp(Group =“Williams“)), alternative=“greater“))
~~~

## 5.4 R-code: Multiple correlated crude tumor incidence using Williams trend tests

~~~
glm24 <− glm(cbind(Successes + 1, 52 - (Successes+1)) ~ Group,
subset(miceN, Site==“t24“), family=binomial())
glm27 <− glm(cbind(Successes + 1, 52 - (Successes+1)) ~ Group,
subset(miceN, Site==“t27“), family=binomial())
glm29 <− glm(cbind(Successes + 1, 52 - (Successes+1)) ~ Group,
subset(miceN, Site==“t29“), family=binomial())
glm30 <− glm(cbind(Successes + 1, 52 - (Successes+1)) ~ Group,
subset(miceN, Site==“t30“), family=binomial())
mTW <−summary(glht(mmm(T24=glm24,T27=glm27,T29=glm29,T30=glm30),
mlf(mcp(Group =“Williams“)), alternative=“greater“))
~~~

## 5.5 R-code: Multiple effect sizes using poly-k=3 Tukey-type trend tests

~~~
miceL$weightpoly24 <− 1
wt024 <− which(miceL$t24 == 0)
miceL$weightpoly24[wt024] <− (miceL$death[wt024]/max(miceL$death))^3
library(glm2)
library(logbin, quietly=TRUE)
t24OR <−glm(t24~dose, data=miceL, family= binomial(link=“logit“), weight=weightpoly24)
t24RD <−glm2(t24~dose, data=miceL, family= binomial(link=“identity“), weight=weightpoly24)
t24RR <−glm2(t24~dose, data=miceL, family= binomial(link=“log“), weight=weightpoly24)
tuOR <− tukeytrendfit(t24OR, dose=“dose”, scaling=c(“ari”, “ord”, “arilog“))
pOR <− summary(glht(model=tuOR$mmm, linfct=tuOR$mlf, alternative=“greater“))
tuRD <− tukeytrendfit(t24RD, dose=“dose”, scaling=c(“ari”, “ord”, “arilog“))
pRD <− summary(glht(model=tuRD$mmm, linfct=tuRD$mlf, alternative=“greater“))
tuRR <− tukeytrendfit(t24RR, dose=“dose”, scaling=c(“ari”, “ord”, “arilog“))
pRR <− summary(glht(model=tuRR$mmm, linfct=tuRR$mlf, alternative=“greater“))
ttany <− combtt(tuOR, tuRD,tuRR)
anylink<−summary(glht(model=ttany$mmm, linfct=ttany$mlf, alternative=“greater“))
~~~

## 5.6 R-code: Pairwise and trend tests jointly: crude tumor incidence of hepatoblastoma

~~~
############### Permutative pairwise and William tests
miceT <−miceF[, c(1, 27), ]# t24 only
miceT$Group <−as.factor(miceT$group) table(miceT)
library(“coin“)
Wh <− c(−1, 0, 0, 1)
Wm <− c(−1, 0, .5,.5)
Wp <− c(−1, 1/3, 1/3, 1/3)
p3 <− c(−1,0,0,1)
p2 <− c(−1,0,1,0)
p1 <− c(−1,1,0,0)
g <− function(x) {
x <− unlist(x)
cbind( W1 = Wh[x], W2 = Wm[x],W3 = Wp[x], P1=p1[x], P2=p2[x])
}
it <− independence_test(t24~ Group, data = miceT, xtrafo = g, alternative = “greater“)
pvalue(it, method = “single-step“)
################ asymptotic Dunnett and Tukey
TWDu5 <− tukeytrendfit(fitpoly5, dose=“dose”,
scaling=c(“ari”, “ord”, “arilog”, “treat“), ctype=“Dunnett“)
twdu5 <− glht(model=TWDu5$mmm, linfct=TW5$mlf, alternative=“greater“)
polytWDu5<−fortify(summary(twdu5))
~~~

## 5.7 R-code: Complete example

~~~
miceZ$weightpoly241 <− 1
miceZ$weightpoly242 <− 1
miceZ$weightpoly243 <− 1
miceZ$weightpoly244 <− 1
miceZ$weightpoly245 <− 1
miceZ$weightpoly246 <− 1
wt024 <− which(miceZ$t24 == 0)
miceZ$weightpoly241[wt024] <− (miceZ$death[wt024]/max(miceZ$death))^1
miceZ$weightpoly242[wt024] <− (miceZ$death[wt024]/max(miceZ$death))^2
miceZ$weightpoly243[wt024] <− (miceZ$death[wt024]/max(miceZ$death))^3
miceZ$weightpoly244[wt024] <− (miceZ$death[wt024]/max(miceZ$death))^4
miceZ$weightpoly245[wt024] <− (miceZ$death[wt024]/max(miceZ$death))^5
miceZ$weightpoly246[wt024] <− (miceZ$death[wt024]/max(miceZ$death))^6
miceZ$weightpoly271 <− 1
miceZ$weightpoly272 <− 1
miceZ$weightpoly273 <− 1
miceZ$weightpoly274 <− 1
miceZ$weightpoly275 <− 1
miceZ$weightpoly276 <− 1
wt027 <− which(miceZ$t27 == 0)
miceZ$weightpoly271[wt027] <− (miceZ$death[wt027]/max(miceZ$death))^1
miceZ$weightpoly272[wt027] <− (miceZ$death[wt027]/max(miceZ$death))^2
miceZ$weightpoly273[wt027] <− (miceZ$death[wt027]/max(miceZ$death))^3
miceZ$weightpoly274[wt027] <− (miceZ$death[wt027]/max(miceZ$death))^4
miceZ$weightpoly275[wt027] <− (miceZ$death[wt027]/max(miceZ$death))^5
miceZ$weightpoly276[wt027] <− (miceZ$death[wt027]/max(miceZ$death))^6
miceZ$weightpoly291 <− 1
miceZ$weightpoly292 <− 1
miceZ$weightpoly293 <− 1
miceZ$weightpoly294 <− 1
miceZ$weightpoly295 <− 1
miceZ$weightpoly296 <− 1
wt029 <− which(miceZ$t29 == 0)
miceZ$weightpoly291[wt029] <− (miceZ$death[wt029]/max(miceZ$death))^1
miceZ$weightpoly292[wt029] <− (miceZ$death[wt029]/max(miceZ$death))^2
miceZ$weightpoly293[wt029] <− (miceZ$death[wt029]/max(miceZ$death))^3
miceZ$weightpoly294[wt029] <− (miceZ$death[wt029]/max(miceZ$death))^4
miceZ$weightpoly295[wt029] <− (miceZ$death[wt029]/max(miceZ$death))^5
miceZ$weightpoly296[wt029] <− (miceZ$death[wt029]/max(miceZ$death))^6
t241OR <−glm(t24~dose, data=miceZ, family= binomial(link=“logit“), weight=weightpoly241)
t241RD <−glm(t24~dose, data=miceZ, family= binomial(link=“identity“), weight=weightpoly241)
t242OR <−glm(t24~dose, data=miceZ, family= binomial(link=“logit“), weight=weightpoly242)
t242RD <−glm(t24~dose, data=miceZ, family= binomial(link=“identity“), weight=weightpoly242)
t243OR <−glm(t24~dose, data=miceZ, family= binomial(link=“logit“), weight=weightpoly243)
t243RD <−glm(t24~dose, data=miceZ, family= binomial(link=“identity“), weight=weightpoly243)
t244OR <−glm(t24~dose, data=miceZ, family= binomial(link=“logit“), weight=weightpoly244)
t244RD <−glm(t24~dose, data=miceZ, family= binomial(link=“identity“), weight=weightpoly244)
t245OR <−glm(t24~dose, data=miceZ, family= binomial(link=“logit“), weight=weightpoly245)
t245RD <−glm(t24~dose, data=miceZ, family= binomial(link=“identity“), weight=weightpoly245)
t246OR <−glm(t24~dose, data=miceZ, family= binomial(link=“logit“), weight=weightpoly246)
t246RD <−glm(t24~dose, data=miceZ, family= binomial(link=“identity“), weight=weightpoly246)
t271OR <−glm(t27~dose, data=miceZ, family= binomial(link=“logit“), weight=weightpoly271)
t272OR <−glm(t27~dose, data=miceZ, family= binomial(link=“logit“), weight=weightpoly272)
t273OR <−glm(t27~dose, data=miceZ, family= binomial(link=“logit“), weight=weightpoly273)
t274OR <−glm(t27~dose, data=miceZ, family= binomial(link=“logit“), weight=weightpoly274)
t275OR <−glm(t27~dose, data=miceZ, family= binomial(link=“logit“), weight=weightpoly275)
t276OR <−glm(t27~dose, data=miceZ, family= binomial(link=“logit“), weight=weightpoly276)
t271RD <−glm(t27~dose, data=miceZ, family= binomial(link=“identity“), weight=weightpoly271)
t272RD <−glm(t27~dose, data=miceZ, family= binomial(link=“identity“), weight=weightpoly272)
t273RD <−glm(t27~dose, data=miceZ, family= binomial(link=“identity“), weight=weightpoly273)
t274RD <−glm(t27~dose, data=miceZ, family= binomial(link=“identity“), weight=weightpoly274)
t275RD <−glm(t27~dose, data=miceZ, family= binomial(link=“identity“), weight=weightpoly275)
t276RD <−glm(t27~dose, data=miceZ, family= binomial(link=“identity“), weight=weightpoly276)
t291OR <−glm(t29~dose, data=miceZ, family= binomial(link=“logit“), weight=weightpoly291)
t292OR <−glm(t29~dose, data=miceZ, family= binomial(link=“logit“), weight=weightpoly292)
t293OR <−glm(t29~dose, data=miceZ, family= binomial(link=“logit“), weight=weightpoly293)
t294OR <−glm(t29~dose, data=miceZ, family= binomial(link=“logit“), weight=weightpoly294)
t295OR <−glm(t29~dose, data=miceZ, family= binomial(link=“logit“), weight=weightpoly295)
t296OR <−glm(t29~dose, data=miceZ, family= binomial(link=“logit“), weight=weightpoly296)
t291RD <−glm(t29~dose, data=miceZ, family= binomial(link=“identity“), weight=weightpoly291)
t292RD <−glm(t29~dose, data=miceZ, family= binomial(link=“identity“), weight=weightpoly292)
t293RD <−glm(t29~dose, data=miceZ, family= binomial(link=“identity“), weight=weightpoly293)
t294RD <−glm(t29~dose, data=miceZ, family= binomial(link=“identity“), weight=weightpoly294)
t295RD <−glm(t29~dose, data=miceZ, family= binomial(link=“identity“), weight=weightpoly295)
t296RD <−glm(t29~dose, data=miceZ, family= binomial(link=“identity“), weight=weightpoly296)
library(tukeytrend)
tuOR241 <− tukeytrendfit(t241OR, dose=“dose”, scaling=c(“ari”, “ord”, “arilog“))
tuOR242 <− tukeytrendfit(t242OR, dose=“dose”, scaling=c(“ari”, “ord”, “arilog“))
tuOR243 <− tukeytrendfit(t243OR, dose=“dose”, scaling=c(“ari”, “ord”, “arilog“))
tuOR244 <− tukeytrendfit(t244OR, dose=“dose”, scaling=c(“ari”, “ord”, “arilog“))
tuOR245 <− tukeytrendfit(t245OR, dose=“dose”, scaling=c(“ari”, “ord”, “arilog“))
tuOR246 <− tukeytrendfit(t246OR, dose=“dose”, scaling=c(“ari”, “ord”, “arilog“))
tuRD241 <− tukeytrendfit(t241RD, dose=“dose”, scaling=c(“ari”, “ord”, “arilog“))
tuRD242 <− tukeytrendfit(t242RD, dose=“dose”, scaling=c(“ari”, “ord”, “arilog“))
tuRD243 <− tukeytrendfit(t243RD, dose=“dose”, scaling=c(“ari”, “ord”, “arilog“))
tuRD244 <− tukeytrendfit(t244RD, dose=“dose”, scaling=c(“ari”, “ord”, “arilog“))
tuRD245 <− tukeytrendfit(t245RD, dose=“dose”, scaling=c(“ari”, “ord”, “arilog“))
tuRD246 <− tukeytrendfit(t246RD, dose=“dose”, scaling=c(“ari”, “ord”, “arilog“))
tuOR271 <− tukeytrendfit(t271OR, dose=“dose”, scaling=c(“ari”, “ord”, “arilog“))
tuOR272 <− tukeytrendfit(t272OR, dose=“dose”, scaling=c(“ari”, “ord”, “arilog“))
tuOR273 <− tukeytrendfit(t273OR, dose=“dose”, scaling=c(“ari”, “ord”, “arilog“))
tuOR274 <− tukeytrendfit(t274OR, dose=“dose”, scaling=c(“ari”, “ord”, “arilog“))
tuOR275 <− tukeytrendfit(t275OR, dose=“dose”, scaling=c(“ari”, “ord”, “arilog“))
tuOR276 <− tukeytrendfit(t276OR, dose=“dose”, scaling=c(“ari”, “ord”, “arilog“))
tuRD271 <− tukeytrendfit(t271RD, dose=“dose”, scaling=c(“ari”, “ord”, “arilog“))
tuRD272 <− tukeytrendfit(t272RD, dose=“dose”, scaling=c(“ari”, “ord”, “arilog“))
tuRD273 <− tukeytrendfit(t273RD, dose=“dose”, scaling=c(“ari”, “ord”, “arilog“))
tuRD274 <− tukeytrendfit(t274RD, dose=“dose”, scaling=c(“ari”, “ord”, “arilog“))
tuRD275 <− tukeytrendfit(t275RD, dose=“dose”, scaling=c(“ari”, “ord”, “arilog“))
tuRD276 <− tukeytrendfit(t276RD, dose=“dose”, scaling=c(“ari”, “ord”, “arilog“))
tuOR291 <− tukeytrendfit(t291OR, dose=“dose”, scaling=c(“ari”, “ord”, “arilog“))
tuOR292 <− tukeytrendfit(t292OR, dose=“dose”, scaling=c(“ari”, “ord”, “arilog“))
tuOR293 <− tukeytrendfit(t293OR, dose=“dose”, scaling=c(“ari”, “ord”, “arilog“))
tuOR294 <− tukeytrendfit(t294OR, dose=“dose”, scaling=c(“ari”, “ord”, “arilog“))
tuOR295 <− tukeytrendfit(t295OR, dose=“dose”, scaling=c(“ari”, “ord”, “arilog“))
tuOR296 <− tukeytrendfit(t296OR, dose=“dose”, scaling=c(“ari”, “ord”, “arilog“))
tuRD291 <− tukeytrendfit(t291RD, dose=“dose”, scaling=c(“ari”, “ord”, “arilog“))
tuRD292 <− tukeytrendfit(t292RD, dose=“dose”, scaling=c(“ari”, “ord”, “arilog“))
tuRD293 <− tukeytrendfit(t293RD, dose=“dose”, scaling=c(“ari”, “ord”, “arilog“))
tuRD294 <− tukeytrendfit(t294RD, dose=“dose”, scaling=c(“ari”, “ord”, “arilog“))
tuRD295 <− tukeytrendfit(t295RD, dose=“dose”, scaling=c(“ari”, “ord”, “arilog“))
tuRD296 <− tukeytrendfit(t296RD, dose=“dose”, scaling=c(“ari”, “ord”, “arilog“))
ttall <− combtt(tuOR241, tuOR242,tuOR243,tuOR244,tuOR245,tuOR246,
tuRD241, tuRD242,tuRD243,tuRD244,tuRD245,tuRD246,
tuOR271,tuOR272,tuOR273,tuOR274,tuOR275,tuOR276,
tuRD271,tuRD272,tuRD273,tuRD274,tuRD275,tuRD276,
tuRD291,tuRD292,tuRD293,tuRD294,tuRD295,tuRD296,
tuOR291,tuOR292,tuOR293,tuOR294,tuOR295,tuOR296)
all<−summary(asglht(ttall))
allU<−summary(asglht(ttall), adjusted(“none“)) library(ggplot2)
ALLU<−fortify(allU) ALL<−fortify(all)
GEM<−cbind(ALL, ALLU$p)
~~~

## 5.8 R-code: Complete example

~~~
ttRD <− combtt(tuRD241, tuRD242,tuRD243,tuRD244,tuRD245,tuRD246,
tuRD271,tuRD272,tuRD273,tuRD274,tuRD275,tuRD276,
tuRD291,tuRD292,tuRD293,tuRD294,tuRD295,tuRD296)
TTRD<−asglht(ttRD)
plot(TTRD, xlab=“Risk difference to p_0”, abline=0, main=“Simultaneous confidence intervals“)
~~~

## 5.9 R-code: Exact tests

~~~
miceZ <−miceF[, c(1:3, 22:38,90)]
miceZ$Group<−as.factor(miceZ$group)
############# Dunnett conditional
library(“coin“)
aa <− c(−1,0,0,1)
dd <− c(−1,0,1,0)
rr <− c(−1,1,0,0)
g <− function(x) {
x <− unlist(x)
cbind( CD1 = rr[x], CD2 = dd[x],CD3 = aa[x])
}
it <− independence_test(t24+t27+t29~ Group, data = miceZ,
xtrafo = g, alternative = “greater“)
pvalM <− round(pvalue(it, method = “single-step“), 14)
################### add1 asymptotic
library(multcomp)
library(arm)
glm24 <− bayesglm(cbind(t24 + .5, 52 - (t27+.5)) ~ Group,data=miceZ,
family=binomial(“identity“))
glm27 <− bayesglm(cbind(t27 + .5, 52 - (t27+.5)) ~ Group,data=miceZ,
family=binomial(“identity“))
glm29 <− bayesglm(cbind(t29 + .5, 52 - (t29+.5)) ~ Group,data=miceZ,
family=binomial(“identity“))
multT <−summary(glht(mmm(T24=glm24, T27=glm27,T29=glm29),
mlf(mcp(Group =“Dunnett“)), alternative=“greater“))
#####multfisher
library(multfisher)
##### pairwise data 0 vs. 1
miceX <−miceF[, c(1,27,30,32)] # crude proportions
MICE03<−droplevels(subset(miceX,group %in% c(0,3)))
MICE03$group[MICE03$group==3]<−1
MICE02<−subset(miceX,group %in% c(0,2))
MICE02$group[MICE02$group==2]<−1
MICE01<−subset(miceX,group %in% c(0,1))
mf01<−mfisher.test(x=MICE01[,c(2,3,4)],y=MICE01$group,method=“alpha”,
closed.test=TRUE,show.region=TRUE, alpha=0.05)
mf02<−mfisher.test(x=MICE02[,c(2,3,4)],y=MICE02$group,method=“number”,
closed.test=TRUE,show.region=TRUE, alpha=0.05)
mf03<−mfisher.test(x=MICE03[,c(2,3,4)],y=MICE03$group,method=“alpha.greedy”,
closed.test=TRUE,show.region=TRUE, alpha=0.05)
round(mf01$elementary.tests$p.adj,digits=19)
round(mf01$closed.test$p.intersect,digits=19)
~~~

